# Structural insights into sphingosine 1-phosphate receptor 4 activation

**DOI:** 10.1101/2024.03.02.583092

**Authors:** Wei Gao, Shiyi Gan, Mengting Zhang, Asuka Inoue, Mengting Xie, Huan He, Huan Zhu, Shanshan Guo, Chen Qiu, Di Chang, Jinling Yu, Zhuo Deng, Fang Ye, Shiliang Li, Jian Zhang, Zhenjiang Zhao, Mengzhu Xue, Bernard Ofosuhene, Yufang Xu, Honghuang Lin, Xuhong Qian, Lili Zhu, Yang Du, Honglin Li

## Abstract

S1PR4 is one of five subtypes of sphingosine 1-phosphate receptors (S1PRs) that regulate immune cell functioning, with functional distinctions to other subtypes. S1PR1-targeted modulators caused serious cardiac and vascular adverse effects because S1PR1 was expressed throughout the whole body. Since S1PR4 was only expressed in lung and lymphoid cells, S1PR4-targeted modulators might not trigger these side effects. However, the development of S1PR4-specific agonists is greatly hindered because of the lack of activated S1PR4 structure. Here, we resolved cryo-EM structures of activated S1PR4 and revealed the structural mechanism of ligand recognition, receptor activation, and Gα_i_ coupling. Our results offered structural templates for the development of selective S1PR4 agonists with improved safety profiles.

## Introduction

Sphingosine 1-phosphate (S1P) is a bioactive lipid mediator originating from ceramide or interconverted with sphingosine, tuning cellular proliferation (*1*), immune cell trafficking (*2-4*), cell motility (*3, 5, 6*), angiogenesis (*3, 7*), vascular maturation (*6, 8, 9*), neurogenesis (*10*), and cardio-protection (*11, 12*). Through its activation of five G-protein coupled S1P receptor subtypes (S1PRs: S1PR1-S1PR5), the S1P-S1PR axis is an effective therapeutic target in several diseases, such as multiple sclerosis (*13*), ulcerative colitis (*14*), and inflammatory bowel disease (*15*). Four S1PR1-targeted modulators (fingolimod (*16*), siponimod (BAF312) (*17*), ozanimod (*18*), and ponesimod (*19*)) are FDA-approved to treat multiple sclerosis. Prior studies have suggested that S1PR4 is essential for regulating the migration of immune cells with functional distinctions to S1PR1 (*20-23*). S1PR4 plays a causative role in various diseases, including psoriasis, steatohepatitis, and mammary tumors (*24-26*).

Although S1PR1, S1PR2, S1PR3, and S1PR5 (*27-33*) structures have been reported, the structure of S1PR4 was unresolved. Thus, it is unknown whether the mechanism of S1PR4 activation distinguishes other S1PRs and leads to functional distinctions from other S1PRs. Here, we resolved cryo-electron microscopy (cryo-EM) structures of apo-S1PR4 and S1P-bound S1PR4 to reveal the structural mechanism of S1PR4 activation.

## Results

### Overall structures of S1PR4-Gα_i_ complex

We resolved the structures of apo- and S1P-bound S1PR4-Gα_i_ through single-particle cryo-EM analysis at resolutions of 3.12Å and 3.42Å, respectively (Fig. 1, A and B; figs. S1 and S2; resolution calculated by model to map; Fourier shell correlation (FSC) = 0.143). We did not observe obvious densities for approximately the first 20 amino acids as well as amino acids 34-49 within S1PR4, suggesting unstable conformations of these regions. The superposition of apo-S1PR4-G_i_ and S1P-S1PR4-G_i_ confirmed that the conformation of the S1PR4 transmembrane domain was very similar (Fig. 1C). The major Gα_i_ binding interface is made up of the polar residues D350 and C351 of the Gα_i_ and the surrounding polar residues R143^3.50^ in TM3 and R79^12.52^ in ICL1 of S1PR4 (Fig. 1E, superscript denotes Ballesteros-Weinstein numbering for GPCRs). However, the G protein binding pattern was changed due to shifting Gα_i_ (Fig. 1, C to E). The C terminus of the α5 helix of Gα_i_ in apo-S1PR4-G_i_ was shifted 2.0Å toward ICL1, causing the N terminus of the α5 helix to move 2.7Å closer towards TM5 than S1P-S1PR4-G_i_ (Fig. 1D, top), resulting in the rotation of αN helix of Gα_i_ (Fig. 1D, bottom). R79^12.52^ forms a salt bridge with the side chain of D350 of Gα_i_ in S1P-S1PR4-G_i_. In contrast, the salt bridge is destroyed, and R79^12.52^ forms a hydrogen bond with the backbone carbonyl of K349 in apo-S1PR4-G_i_. Moreover, K245^6.29^ forms a hydrogen bond with F354, while R239^ICL3^ forms two hydrogen bonds with I319 in the S1P-bound state (Fig. 1E, right), and these interactions are absent in the apo state (Fig. 1E, left). Our findings indicate that the interaction between S1PR4 and Gα_i_ is more stable in the presence of S1P.

**Fig. 1.**
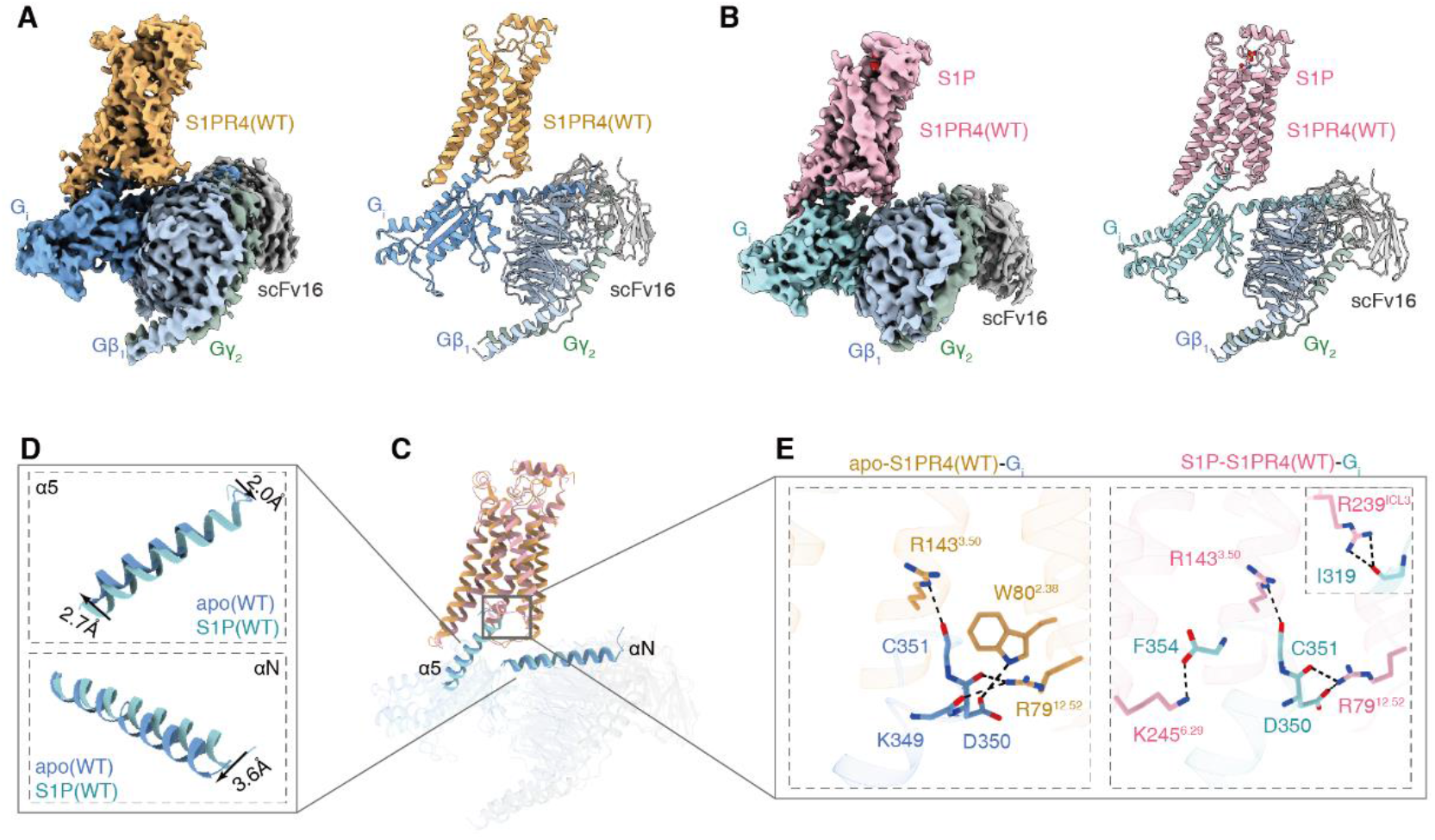
Overall structures of S1PR4-Gα_i_ complex. (**A** and **B**) Orthogonal displays of the cryo-EM density map (left) and ribbon representation (right) of the apo-S1PR4(WT)-G_i_ complex (A) and the S1P-S1PR4(WT)-G_i_ complex (B). The maps are colored based on different subunits. (**C**) Superposition of the G-protein coupling interfaces of the apo-S1PR4(WT)-G_i_ complex and the S1P-S1PR4(WT)-G_i_ complex using the receptor for alignment. (**D** and **E**) Details of α5 and αN helices of Gα_i_ (D) and S1PR4-Gα_i_ binding interface (E) in apo-S1PR4(WT) and S1P-S1PR4(WT). The polar interactions are delineated by black dashed lines.

## Discussion

Since S1PR1 was expressed in many organs, including the brain, heart, spleen, liver, lung, thymus, kidney, skeletal muscle, and lymphoid cells, S1PR1-targeted modulators, such as fingolimod, caused serious cardiac and vascular adverse effects, including hypertension, macular oedema, bradycardic, and atrioventricular block (*34*). In contrast, S1PR4 was only expressed in lung and lymphoid cells. Therefore, S1PR4-targeted modulators might not trigger these side effects. In addition, S1PR4 is essential for promoting Th17 polarization, so it might be a promising target for Th17-associated diseases, including psoriasis, inflammatory bowel disease, and asthma (*20*). However, the structure of S1PR4 bound to agonists was not resolved yet, limiting the development of selective S1PR4 drugs. We resolved cryo-EM structures of activated S1PR4 and revealed the structural mechanism of ligand recognition, receptor activation, and Gα_i_ coupling. Our results offered structural templates for the development of selective S1PR4 agonists for the treatment of Th17-associated diseases with improved safety profiles.

### Materials and Methods Constructs

For the protein expression, human S1PR4 was cloned into a pFastBac vector with the HA signal sequence and a flag epitope tag. To increase the expression level of S1PR4, a BRIL was inserted at the N-terminal of S1PR4. The dominant-negative human Gα_i1_ mutant was cloned into the pFastBac vector. The human Gβ_1_ and the human Gγ_2_ were cloned into a pFastBac-dual vector.

### Purification of S1PR4-DNG_i_ complex

Sf9 insect cells were co-infected with S1PR4, Gα_i1_ and Gβ_1_γ_2_ viruses and cultured for 48 hours. Cell pellets were lyzed in a buffer containing 10 mM Tris pH 7.5, 1 mM EDTA and 1 mg/mL iodoacetamide. Then, cells were resuspended in solubilization buffer containing 20 mM HEPES pH 7.5, 100 mM NaCl, 10% Glycerol, 1% n-Dodecyl-B-D-Maltoside (DDM), 0.1% cholesteryl hemisuccinate (CHS), 160 μg/mL benzamidine, 100 μg/mL leupeptin, 1 mg/mL iodoacetamide, apyrase (25 mU/mL), 10 mM MgCl_2_ and 10 mM CaCl_2_ supplemented with agonist. The suspension was incubated for 1 h at 4 °C. The supernatant was collected by centrifugation and incubated with M1 anti-Flag affinity resin. DDM was exchanged by lauryl maltose neopentyl glycol (LMNG). The S1PR4-DNG_i_ complex was eluted with elution buffer containing 5 mM EDTA and 0.2 mg/mL flag peptide. The eluted complex was incubated with scFv16 for 2 h at 4 °C followed by purification on a Superdex 200 10/300 increase size exclusion column.

### Cryo-EM sample preparation and data collection

The gold film (*35*) (UltraAuFoil/QuantiFoil, 300 mesh, R1.2/1.3, Quantifoil Micro Tools GmbH, DE) or amorphous alloy film (*36*) (300 mesh, R1.2/1.3, Zhenjiang Lehua Electronic Technology Co., Ltd.) was glow discharged with air for 40 s at 15 mA at easiGlow™ Glow Discharge Cleaning System (PELCO, USA) or Tergeo Plasma Cleaner (PIE SCIENTIFIC, USA). 3 μL purified complex sample was dropped onto the grid and then blotted for 3-4 s with blotting force 0 and plunged into liquid ethane cooled by liquid nitrogen using Vitrobot Mark IV (Thermo Fisher Scientific, USA). Cryo-EM datasets were collected with the 300 kV Titan Krios Gi3 microscope. The raw movies were collected by Gatan K3 BioQuantum Camera at the magnification of 105000, with a pixel size of 0.85 Å. Inelastically scattered electrons were excluded by a GIF Quantum energy filter (Gatan, USA) using a slit width of 20 eV. The movies were acquired with the defocus range of -1.2 to -2.0 μm with a total exposure time of 2 s fragmented into 50 frames and with a dose rate around 15.36 e/pixel/s. SerialEM was used for semi-automatic data acquisition (*37*).

### Cryo-EM data processing, model building and refinement

The image stacks were collected and subjected to motion correction using MotionCor2 (*38*). Contrast transfer function parameters were estimated by CTFFIND4 (*39*), implemented in RELION3.1/4.0 (*40*) or Patch CTF in cryoSPARC (*41*). Particles were auto-picked from micrographs by RELION/cryoSPARC and then subjected to 2D classification using cryoSPARC. Selected particles with an appropriate 2D average from 2D classification were further subjected to 3D classification using cryoSPARC. Eventually, particles with high-resolution 3D average were selected from 3D classification, which resulted in a map with an initial resolution of near-atomic level by NU-refinement. The refined particles were subjected to CTF refinement to update per-particle defocus and per-micrograph astigmatism. Followed by sharpening in cryoSPARC, a series of final maps with resolution from 3.12 Å to 3.42 Å were generated after postprocessing determined by gold-standard Fourier shell correlation using the 0.143 criteria. The local resolution map was calculated from cryoSPARC using two unfiltered half maps. The predicted human-S1PR4 structure obtained from Alphafold (https://alphafold.ebi.ac.uk/) was used as a template to build the S1PR4-G_i_-scFv16 complex model. The template for G_i_-scFV16 was obtained from the G_i_ heterotrimer of S1PR1-G_i_ cryoEM structure (PDB 7EO2). Model docking was carried out using Chimera (*42*). Manually adjustment and rebuilding was performed with COOT (*43*). The model was repeatedly refined using real space refinement module in Phenix (*44*). The softwares, such as UCSF Chimera and UCSF ChimeraX (*45*), were used to prepare the molecular graphics figures. The final model statistical analysis was performed by Phenix.

## Data availability

The density maps and structure coordinates have been deposited in the Electron Microscopy Data Bank (EMDB) and the Protein Data Bank (PDB) with accession numbers EMD-37783 and 8WRN, respectively, for the apo-S1PR4-G_i_ complex; and EMD-37817 and 8WSL, respectively, for the S1P-S1PR4-G_i_ complex.

## Acknowledgments

We would like to thank the Kobilka Cryo-Electron Microscopy Center, the Chinese University of Hong Kong, Shenzhen for our cryo-EM study. This study was supported in part by the National Natural Science Foundation of China (grants 81825020 and 82150208 to H. Li; 32271263 to Y. Du), the National Key R&D Program of China (grants 2022YFC3400501 and 2022YFC3400504 to H. Li), the Sci & Tech Innovation Commission of Shenzhen (JCYG 20220818103009018 to Y. Du). Honglin Li is also sponsored by the National Program for Special Supports of Eminent Professionals and National Program for Support of Top-notch Young Professionals. Yang Du is also sponsored by the Ganghong Young Scholar Development Fund, and in part by the Kobilka Institute of Innovative Drug Discovery at the Chinese University of Hong Kong, Shenzhen. Asuka Inoue was funded by KAKENHI JP21H04791 and JP21H05113 from Japan by the Society for the Promotion of Science (JSPS), JPMJFR215T and JPMJMS2023 from Japan Science and Technology Agency (JST), JP22ama121038 and JP22zf0127007 from the Japan Agency for Medical Research and Development (AMED).

## Author contributions

Data acquisition and progression: W. Gao, S. Gan, M. Zhang, A. Inoue, M. Xie, H. He, H. Zhu, S. Guo, C. Qiu, D. Chang, J. Yu, Z. Deng, and F. Ye. Data interpretation: W. Gao, S. Li, Z. Zhao, Y. Xu, and X. Qian. Manuscript draft: W. Gao, S. Gan, M. Zhang, and L. Zhu. Manuscript editing and review: J. Zhang, M. Xue, B. Ofosuhene, H. Lin, L. Zhu, Y. Du, and H. Li. Conceptualization: H. Li, Y. Du, and L. Zhu.

---

**Fig. S1.**
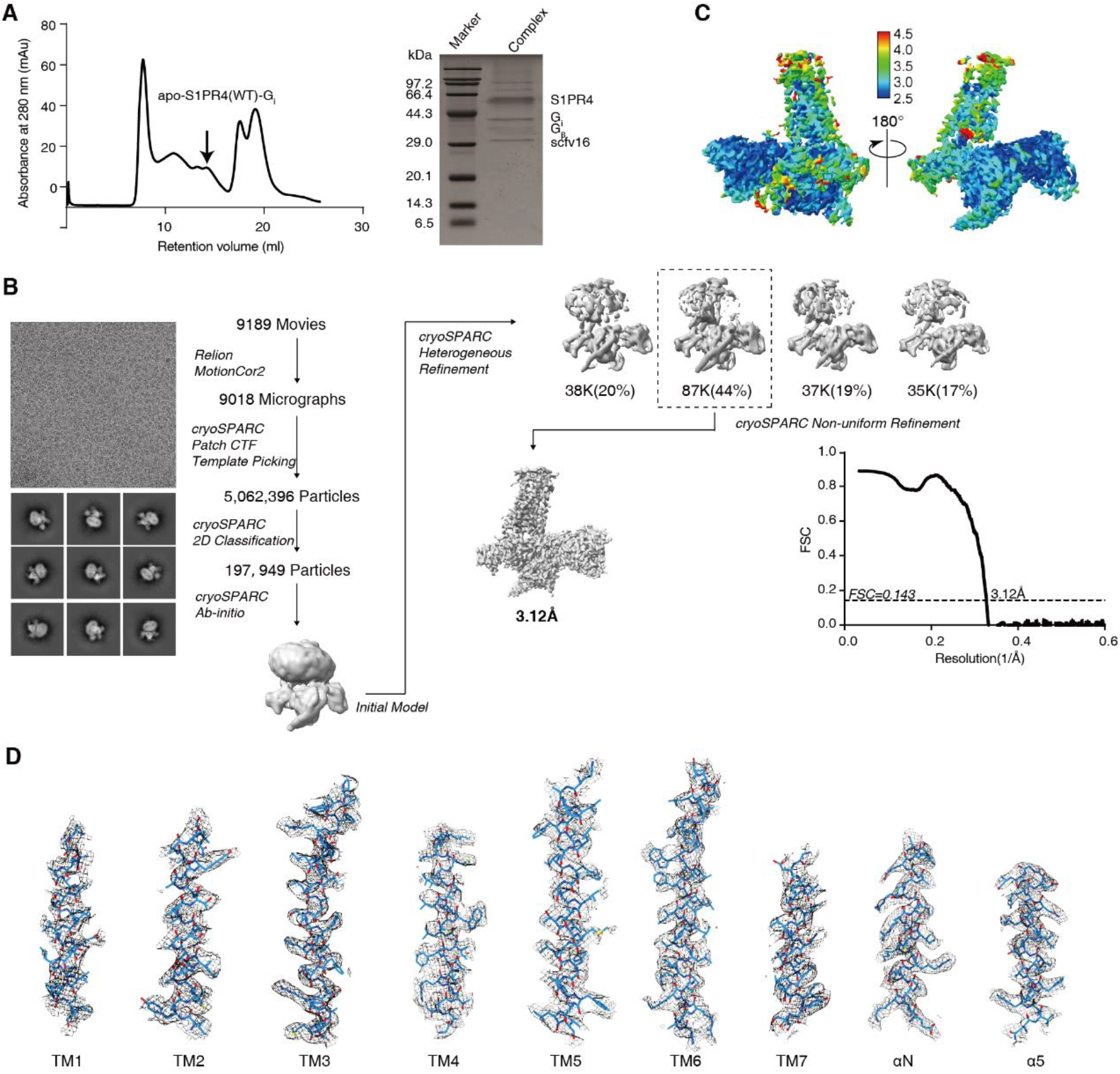
Single-particle reconstruction of the apo-S1PR4(WT)-G_i_ complex. (**A**) Size-exclusion chromatography and SDS-PAGE of the apo-S1PR4(WT)-G_i_ complex. (**B**) Cryo-EM micrograph, 2D class averages, and flow chart of cryo-EM analysis for the apo-S1PR4(WT)-G_i_ complex. (**C**) Local resolution display of the reconstructed 3.12 Å map. A local resolution of 3.12 Å map was projected within cryoSPARC. The resolution range is depicted from 2.5 Å-4.5 Å. (**D**) Side-chain density of the TM helix of S1PR4 as well as Gα_i_ αN and Gα_i_ α5 helices of the 3.12 Å map. Volume was contoured at a threshold level of 0.300.

**Fig. S2.**
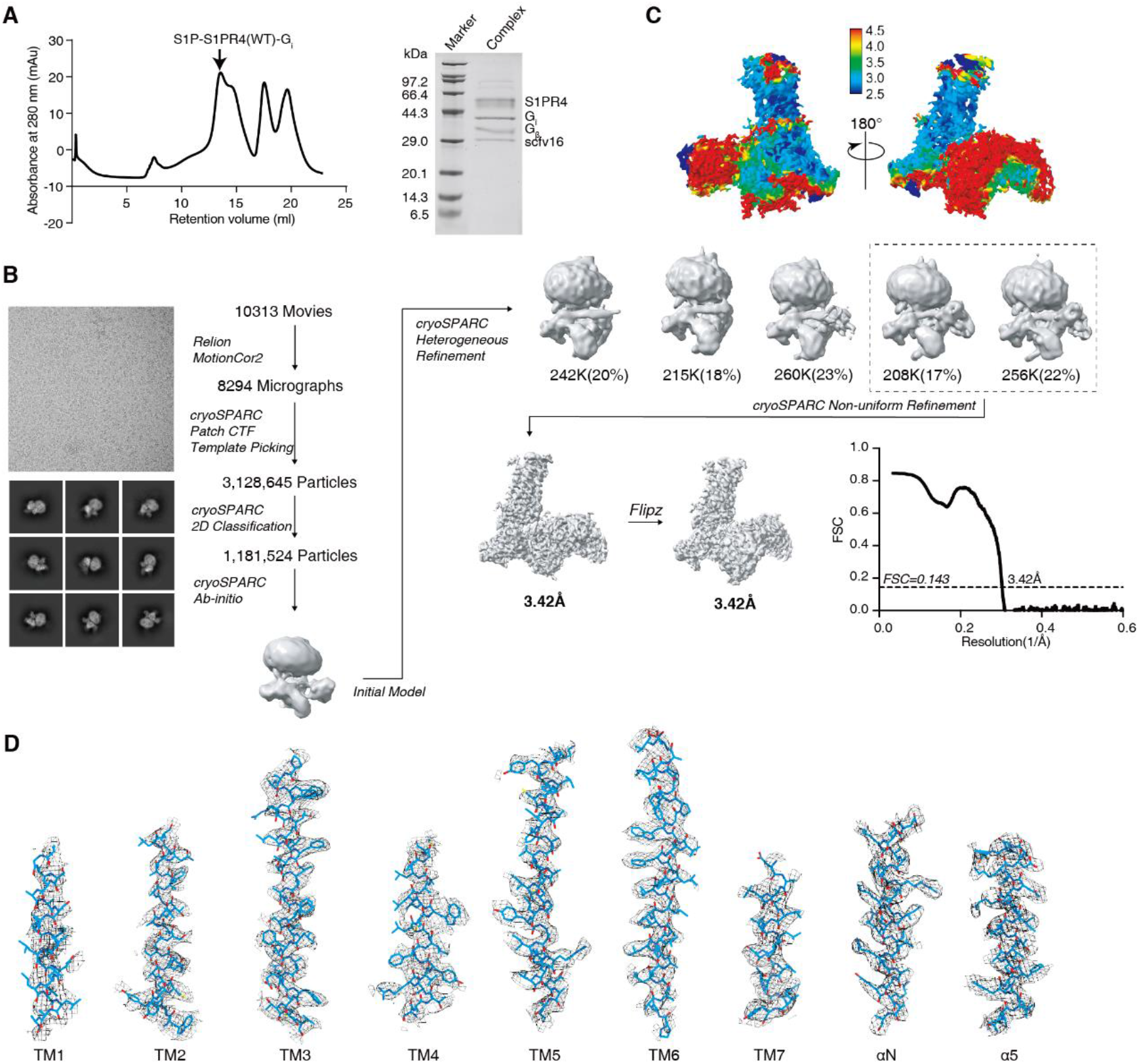
Single-particle reconstruction of the S1P-S1PR4(WT)-G_i_ complex. (**A**) Size-exclusion chromatography and SDS-PAGE of the S1P-S1PR4(WT)-G_i_ complex. (**B**) Cryo-EM micrograph, 2D class averages, and flow chart of cryo-EM analysis for the S1P-S1PR4(WT)-G_i_ complex. (**C**) Local resolution display of the reconstructed 3.42 Å map. A local resolution of 3.42 Å map was projected within cryoSPARC. The resolution range is depicted from 2.5 Å-4.5 Å. (**D**) Side-chain density of TM helix of S1PR4 as well as Gα_i_ αN and Gα_i_ α5 helices of the 3.42 Å map. Volume was contoured at a threshold level of 0.303.

